# *N*-aldehyde modified phosphatidylethanolamines formed by lipid peroxidation inhibit efferocytosis

**DOI:** 10.64898/2026.09.10.750792

**Authors:** Reza Fadaei, Annie Bernstein, Elizabeth A. Wayman, Azuah L. Gonzalez, Keri A. Tallman, Zahra Mashhadi, Abdul-musawwir Alli-oluwafuyi, Sergey Dikalov, MacRae F. Linton, Kasey C. Vickers, Amanda C. Doran, Sean S. Davies

## Abstract

**Background:** Impairments in efferocytosis, the phagocytic clearance of apoptotic cells, contribute to the development of chronic inflammation and promote the formation of large necrotic cores within atherosclerotic plaques. A better understanding of the factors that impair efferocytosis could lead to improved interventions. Many chronic inflammatory conditions feature excessive lipid peroxidation, which generates reactive lipid aldehydes that can potentially form *N*-aldehyde-modified phosphatidylethanolamines (NAMPs). We therefore investigated the extent to which levels of NAMPs are elevated under conditions of oxidative stress and whether exposure of macrophages to NAMPs inhibit efferocytosis.

**Methods:** The effects of synthetic NAMPs and lipid peroxidation on efferocytosis were measured using the Incucyte Live-Cell Analysis System to monitor rate of ingestion of CypHer-labeled apoptotic cells by cultured bone marrow-derived macrophages in real time. The extent of NAMP formation under various conditions was examined using liquid chromatography coupled with mass spectrometry.

**Results:** Several representative species of NAMPs (*N*-isolevuglandin-phosphatidylethanolamine, *N*-IsoLG-PE; *N*-4-hydroxynonenal-phosphatidylethanolamine, *N*-HNE-PE; and *N*-azeloyl-phosphatidylethanolamine, N-Aze-PE) inhibited efferocytosis in a concentration-dependent manner. Hydrolysis of NAMPs by recombinant *N*-acyl phosphatidylethanolamine hydrolyzing phospholipase D (NAPE-PLD) abolished their inhibitory activity, whereas genetic deletion of macrophage NAPE-PLD enhanced NAMP-mediated inhibition of efferocytosis. Blocking the inhibitor of lipid peroxidation, glutathione peroxidase 4 (GPX4), increased macrophage NAMP concentrations and impaired efferocytosis. Unmodified high-density lipoprotein (HDL) promoted efferocytosis, but exposing HDL to reactive lipid aldehydes or peroxidizing agents markedly increased NAMPs and this modified HDL inhibited efferocytosis. HDL isolated from subjects with familial hypercholesterolemia had elevated levels of multiple NAMP species. NAMPs impaired macrophage cholesterol efflux, which is required for continual efferocytosis.

**Conclusions:** NAMPs accumulate under conditions associated with lipid peroxidation and inhibit the ability of macrophages to carry out efferocytosis.

**What are the clinical implications of this study?:** Our finding that NAMPs inhibit efferocytosis by macrophages suggests that interventions that reduce NAMP formation such as lipid aldehyde scavenging or increasing NAPE-PLD expression or activity to enhance NAMP degradation represent novel therapeutic strategies to restore efferocytosis in atherosclerosis and other chronic inflammatory diseases.

## INTRODUCTION

Although inflammatory responses are critical to host defense against pathogens and traumatic injury, the inability to appropriately constrain and resolve inflammation leads to the development of chronic inflammatory diseases such as atherosclerotic cardiovascular disease (ASCVD). A key step in the resolution of inflammation is the efficient phagocytic clearance of apoptotic immune cells that accumulate during the initial inflammatory response (i.e. efferocytosis)^1, 2^. Absent such clearance, apoptotic cells undergo secondary necrosis, releasing their intracellular content of proteases, nucleases, and lipases, resulting in damage to nearby cells and creating damage-associated pattern recognition molecules that prolong inflammatory responses^2^. Multiple lines of evidence implicate impaired efferocytosis in chronic inflammatory diseases^3^. In ASCVD, impaired efferocytosis drives the formation and expansion of necrotic cores within atherosclerotic lesions, a hallmark of plaques vulnerable to rupture^1, 2^ ^4^.

Peroxidized lipids and lipoproteins form during atherogenesis and drive inflammatory processes^5–7^; however, whether they also impair the resolution of inflammation has not yet been established. Peroxidation of polyunsaturated fatty acids generates a variety of reactive lipid aldehydes including isolevuglandins (IsoLG), 4-hydroxynonenal (HNE), 4-oxo-nonenal (ONE), 9-keto-12-oxo-10-dodecenoic acid (KODA), and 9-oxononanoic acid^8–13^ (**Fig. 1A**). These lipid aldehydes covalently modify primary amines including phosphatidylethanolamine, thereby generating a series of *N*-aldehyde modified phosphatidylethanolamines (NAMPs)^14^ ^15, 16^ (**Fig. 1B**). Several NAMP species including *N*-IsoLG-PE and *N-*HNE-PE are potent inducers of inflammation^14^, suggesting they act as damage-associated molecular pattern molecules. *N*-IsoLG-PE signals through the receptor for advanced glycation endproducts (RAGE) to induce Nuclear Factor Kappa B (NFkB) activation^17^. *N*-IsoLG-PE also induces ER stress^18^ and inhibits the function of complex I of the mitochondrial oxidative phosphorylation complex^19^, which can also promote inflammation.

**Figure 1.**
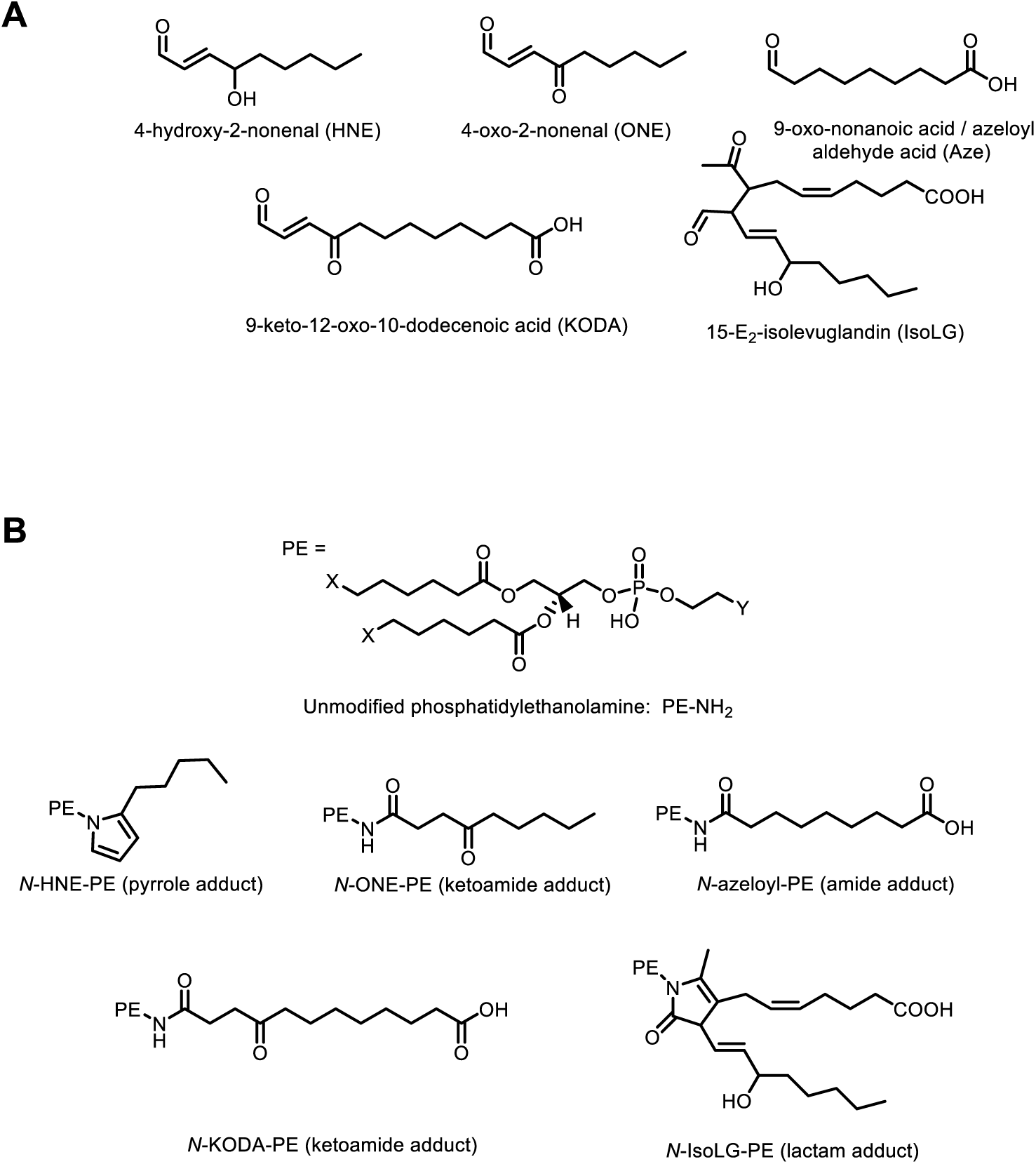
Chemical structures. The chemical structures of (A) reactive lipid aldehydes and (B) their corresponding *N*-aldehyde-modified phosphatidylethanolamine (NAMP). For each aldehyde, the predominant NAMP species is shown.

*N*-acyl phosphatidylethanolamine phospholipase D (NAPE-PLD) is an evolutionarily conserved enzyme whose expression is reduced in atherosclerotic lesions^20^. Bone marrow–derived macrophages (BMDMs) from *Napepld^-/-^*(KO) mice show reduced ability to carry out efferocytosis than BMDMs from *Napepld^+/+^* (WT) mice^21^, supporting a role for NAPE-PLD in facilitating efferocytosis. NAPE-PLD catalyzes two reactions that may be necessary to promote efficient efferocytosis: the hydrolysis of *N*-acyl-phosphatidylethanolamines (NAPEs)^22, 23^ to produce pro-efferocytic fatty acid ethanolamides (FAEs) such as palmitoylethanolamide (PEA) and oleoylethanolamide (OEA) and the hydrolysis of NAMPs^16^, thereby potentially inactivating them. We hypothesized that NAMPs inhibit efferocytosis, so that conditions that cause NAMP accumulation, including reductions in NAPE-PLD activity, would cause impaired efferocytosis. We therefore examined the effect of NAMPs on the ability of cultured macrophages to carry out efferocytosis and whether conditions that increase NAMP accumulation also impair efferocytosis.

## MATERIALS and METHODS

### Materials

Materials used for cell culture were from Gibco by Life Technologies, Inc. *N*-6,2′-O-dibutyryladenosine 3′,5′-cyclic monophosphate (cAMP), methyl-β-cyclodextrin, triethylammonium acetate (TEAA), Sandoz 58-035, FPS-ZM1, and BLT-1 were purchased from Sigma-Aldrich. CypHer5E NHS ester was obtained from Cytiva. TopFluor® cholesterol (BODIPY-cholesterol) and dipalmitoyl phosphatidylethanolamine (dpPE 16:0) were purchased were purchased from Avanti Polar Lipids. 4-hydroxy-2-nonenal (HNE) and 4-oxo-2-nonenal (ONE) were obtained from Cayman Chemical. TAK-242 was obtained from Tocris Bioscience. V70 (azobis[4-methoxy-2,4-dimethyl valeronitrile]) was purchased from Wako Chemicals. 9-(tert-butoxy)-9-oxononanoic acid was purchased from Combi-Blocks. Organic solvents were LCMS grade from EMD Millipore. HDL obtained from fasting subjects was isolated by density gradient ultracentrifugation and dialyzed into PBS.

### Recombinant NAPE-PLD

Recombinant NAPE-PLD was expressed and purified using nickel-affinity chromatography with minor modifications to previously described methods^16^. Briefly, bacterial pellets were lysed in His-binding buffer (50 mM Tris-HCl, 0.5 M NaCl, 5 mM imidazole, pH 7.8) supplemented with protease inhibitors and 0.1% Triton X-100, followed by clarification by centrifugation. The supernatant was applied to Ni-NTA resin, washed with wash buffer (50 mM Tris-HCl, 0.5 M NaCl, 60 mM imidazole, pH 7.8), including an additional wash containing 0.3 M guanidine to reduce non-specific binding, and eluted using elution buffer (50 mM Tris-HCl, 0.5 M NaCl, 0.5 M imidazole, pH 7.8). Fractions containing NAPE-PLD were identified by SDS-PAGE.

### Synthesis of NAMPs

*N*-IsoLG-PE, *N*-HNE-PE, *N*-ONE-PE, and *N*-KODA-PE were synthesized by overnight incubation of 500 nM dipalmitoyl phosphatidylethanolamine with 2, 10, 3, and 5 molar equivalents of IsoLG, HNE, ONE, and KODA, respectively, in triethylammonium acetate/isopropanol buffer (1:1, v/v), as previously described^16^. Reaction products were extracted with chloroform/methanol, and generation of NAMP species was verified and quantified by LC–MS. *N*-Aze-PE was synthesized by reaction of 9-(tert-butoxy)-9-oxononanoic acid with dipalmitoyl PE in the presence of coupling reagents (**Supplemental Figure 1**), with subsequent removal of the protecting group generating a product whose 1H-NMR spectrum was consistent with *N*-Aze-PE (**Supplemental Figure 2**).

### Oxidation of HDL

HDL was peroxidized by incubation of 300 µL of HDL (12 mg protein/mL) with V70 (1 mM) at 37 °C overnight under oxygen-rich conditions. Unmodified HDL was prepared using the same condition without V70 and oxygen-rich conditions. For experiments where 2-hydroxybenzylamine (2HOBA) was added, 2HOBA (100 mM) was added prior to the addition of V70. For experiments where recombinant NAPE-PLD was added, after the overnight incubation with V70, active and heat inactivated NAPE-PLD (100 ng/mL) were added to the samples for an overnight incubation in 37 °C. Aliquots of unmodified and peroxidized HDL were stored at −80 °C until use.

### Measurement of NAMPs

Lipids were extracted from samples using chloroform/methanol/water (8:4:3, v/v/v) and dried under nitrogen, then subjected to methylamine hydrolysis by incubating with methanol/methylamine/1-butanol (4:4:1, v/v/v) at 53 °C for 1 hour to selectively remove the O-acyl chains while preserving the shared *N*-aldehyde-modified ethanolamine glycerophosphate of each NAMP species. This treatment enhances analytical sensitivity for NAMP species. Samples were subsequently dried under nitrogen gas, reconstituted in mobile phase B, and analyzed by LC–MS as previously described^16^. Briefly, samples were analyzed using a Thermo Q-Exactive Orbitrap mass spectrometer coupled to a Dionex UPLC system (Thermo, San Jose, CA). Chromatographic separation was performed on a Kinetex C8 column (2.6 µm, 100 Å; 100 × 2.1 mm), with the column temperature maintained at 40 °C. The flow rate was set to 0.25 mL/min and the injection volume was 5 µL. Samples were reconstituted in mobile phase B prior to injection. Mobile phase A consisted of 90% water and 10% acetonitrile containing 15 mM ammonium formate and 0.2% (v/v) formic acid. Mobile phase B consisted of acetonitrile/methanol/water (90/5/5 v/v/v) with 0.2% (v/v) formic acid. The gradient program was as follows: 0–1.5 min, 2% B (hold); 1.5–5 min, 2% to 30% B; 5–8 min, 30% to 98% B; 8–9 min, 98% B; 9–9.2 min, 98% to 2% B; and 9.2–10 min, 2% B. Mass spectrometry was performed in negative ion mode over an m/z range of 100–750. Electrospray ionization was carried out using a heated ESI source with a spray voltage of 3,500 V, sheath gas set to 40, capillary temperature maintained at 320 °C, and S-lens RF level set to 50. The scan rate was 6 Hz. Data were acquired and processed using Thermo Xcalibur Qual Browser software. Reconstructed single-ion monitoring chromatograms were generated by setting the plot type to mass range and specifying the calculated exact mass to four decimal places, using a 13-point Gaussian smoothing function and a mass tolerance of 3 ppm. Identification of NAMP species was confirmed by comparison with synthetic NAMP standards analyzed in the same matrix, with identity assigned based on matching exact mass and retention time.

### Derivation of Bone Marrow-Derived Macrophages (BMDM)

For studies involving only WT BMDM, wild-type C57BL/6J mice were obtained from Jackson Lab (stock #0064). For studies comparing WT vs KO BMDM, *Napepld^-/+^* mice were bred together and the resulting *Napepld^+/+^* (WT) or *Napepld^-/-^* (KO) littermates used to generate BMDM. The original *Napepld^-/-^* mouse strain was generated by Drs. Palmiter and Luquet^24^ and provided to Dr. Ken Mackie at Indiana University^25^, with breeding pairs from the Indiana University colony used to establish the Vanderbilt colony. Genotyping of mice in the Vanderbilt colony was performed by MyColony. Mice were sacrificed by isoflurane anesthesia, after which femurs and tibias were excised. Bone marrow cells were collected by flushing the bones with DMEM (4.5 g/L glucose) using a 26-gauge needle. The resulting cell suspension was filtered through a 70-µm cell strainer, pelleted by centrifugation at 500 × g for 5 minutes, and resuspended in macrophage differentiation medium consisting of DMEM (4.5 g/L glucose) supplemented with 20% L-929–conditioned medium, 10% heat-inactivated fetal bovine serum, and 1% penicillin/streptomycin. 25% of the cells were distributed in macrophage differentiation medium 10 mL per plate into a 100-mm culture dish and incubated at 37 °C in a humidified 5% CO₂ atmosphere. After 4 days, nonadherent cells and debris were removed and replaced with fresh differentiation medium. Macrophages were allowed to differentiate for a total of 7 days, with media changes every 2–3 days, before being collected for in vitro efferocytosis or other studies.

### *In vitro* efferocytosis assays

Bone marrow–derived macrophages were plated in 96-well culture plates at a density of 10 × 10^3^ cells per well and allowed to attach overnight. Jurkat cells were rendered apoptotic by exposure to ultraviolet irradiation (254 nm) for 5 min, followed by incubation at 37 °C in a 5% CO₂ environment for 2 h. Using this protocol, routine quality control staining demonstrated that >85% of cells were in early apoptosis, as indicated by annexin V positivity and propidium iodide exclusion. Apoptotic Jurkat cells were fluorescently labeled with pH-sensitive CypHer5E NHS Ester (Cytivia) with 1 x10^6^ cells/mL concentration according to the manufacturer’s protocol and subsequently resuspended in macrophage culture medium at 0.2 × 10^6^ cells/mL. After treatment of BMDM with various compounds (see below), labeled apoptotic Jurkat cells were added at a 2:1 ratio of apoptotic cells to macrophages. Upon engulfment into the acidic environment of macrophage phagolysosomes, CypHer fluorescence increases, thereby providing a real-time measure of efferocytic activity. Phase-contrast and red fluorescence images were acquired for 4 hours using the IncuCyte® S3 Live-Cell Analysis System (Sartorius) imaging platform and images analyzed using the Incucyte integrated image analysis software. Efferocytosis was quantified by measuring the total integrated red fluorescence intensity per image, which served as an indicator of phagocytosed target cells and was used to compare efferocytosis across experimental conditions.

### Cultured macrophage treatment studies

BMDMs were treated with NAMPs, HDL, or oxHDL for 6 h prior to the addition of apoptotic Jurkat cells. In experiments using antagonists or inhibitors, cells were pretreated for 30 minutes before the 6 h treatment period. Hydrolysis of NAMPs by NAPE-PLD was performed as previously described^16^. Briefly, reaction mixtures containing 15 µM NAMPs, 0.4% N-octyl glucoside, 50 mM Tris-HCl, and 20 ng/mL NAPE-PLD were incubated at 37°C for 2 h. Lipids were then extracted using 8:4:3 chloroform/methanol/water. The organic phase was dried under nitrogen and reconstituted in DMSO for treatment of BMDMs. Control samples containing heat-inactivated (95°C for 10 min) enzyme were processed in parallel. The maximum concentration of NAMPs or RSL3 chosen for testing was based on the maximum concentration that showed no detectable cytotoxicity after 18 h of treatment as assessed by the MTT viability assay (not shown).

### Superoxide measurements in NAMP-treated macrophages

Confluent BMDMs in 100-mm dishes were treated with 3 µM *N*-Aze-PE and 1 µM *N*-IsoLG-PE for 6 h in DMEM cell culture. Cells were washed twice with Krebs-Hepes buffer (containing 5.786 g/L NaCl, 0.35 g/L KCl, 0.368 g/L CaCl2, 0.296 g/L MgSO4, 2.1 g/L NaHCO3, 0.142 g/L K2HPO4, 5.206 g/L Na-Hepes, 2 g/L D-glucose, pH = 7.35) and then incubated with 0.5 mM spin probe TMH (Enzo, ALX-430-132) in Krebs-Hepes buffer supplemented with 0.1 mM DTPA for 60 minutes at 37°C as previously described^26^. Supernatant was removed, cells were collected and placed into a 1 ml syringe with 0.6 ml Krebs-Hepes buffer plus DTPA and the suspension was snap-frozen in liquid nitrogen. Samples were analyzed in finger Dewar vessel filled with liquid nitrogen^27^. Total cellular superoxide production was determined by accumulation of superoxide-mediated oxidation of TMH into TM-nitroxide using Electron Spin Resonance Bruker EMX plus spectrometer. Spectrometer settings were as follows: field sweep, 160 G; microwave frequency, 9.43 GHz; microwave power, 2 mW; modulation amplitude, 5 G; conversion time, 36.7 ms; time constant 5.2 s; sweep time, 300 s.

### Cholesterol efflux studies

Cholesterol efflux was measured using a BODIPY-cholesterol–based assay performed as previously described^28^ with minor modifications. Cells were seeded at 7 × 10⁴ cells/well and labeled for 1 h at 37 °C in DMEM containing 1% FBS, 1% penicillin/streptomycin, 2 µg/mL ACAT inhibitor, and BODIPY-cholesterol/cholesterol complexed with cyclodextrin at a final BODIPY-cholesterol concentration of 25 µM. Following labeling, cells were equilibrated overnight in serum-free DMEM supplemented with 2 µg/mL ACAT inhibitor, 0.3 mM cAMP to induce ABCA1 expression, and 2 mg/mL BSA as a carrier. Efflux was initiated by replacing the media with macrophage media containing 10% FBS and NAMPs, followed by incubation for 6 h at 37 °C. Fluorescence in collected supernatants and corresponding cell lysates was measured using a plate reader (Ex: 485 nm, Em: 515 nm). Cholesterol efflux was calculated as the percentage of fluorescence in the supernatant relative to total fluorescence (supernatant + cell lysate).

### Statistical Analysis

Statistical analysis was performed using GraphPad Prism version 10.6.1. For studies with concentration response curves or with multiple stimuli, statistical analysis was performed using ordinary 1-way ANOVA followed by Dunnett’s multiple-comparisons test, with data presented as mean and SEM. For studies comparing two conditions where data was normally distributed, an unpaired two-tailed t-test was used and when data was non-normally distributed, groups were compared using Mann-Whitney test.

## RESULTS

Five representative NAMP species were synthesized to examine the effect of NAMPs on efferocytosis: *N-*IsoLG-PE, *N*-HNE-PE, *N*-Aze-PE, *N*-ONE-PE, and *N*-KODA-PE (**Fig. 1B**). BMDM pre-treated with *N*-IsoLG-PE (**Fig. 2A**), *N*-HNE-PE (**Fig. 2B**), or *N*-Aze-PE (**Fig. 2C**) showed markedly reduced ability to carry out efferocytosis, as measured by their ability to bind and internalize labeled apoptotic cells, compared to BMDM pre-treated with vehicle only (DMSO), with the inhibitory effect being concentration-dependent. BMDM pretreated with *N*-ONE-PE (**Fig. 2D**) and *N*-KODA-PE (**Fig. 2E**) showed only insignificant reductions in efferocytosis.

**Figure 2.**
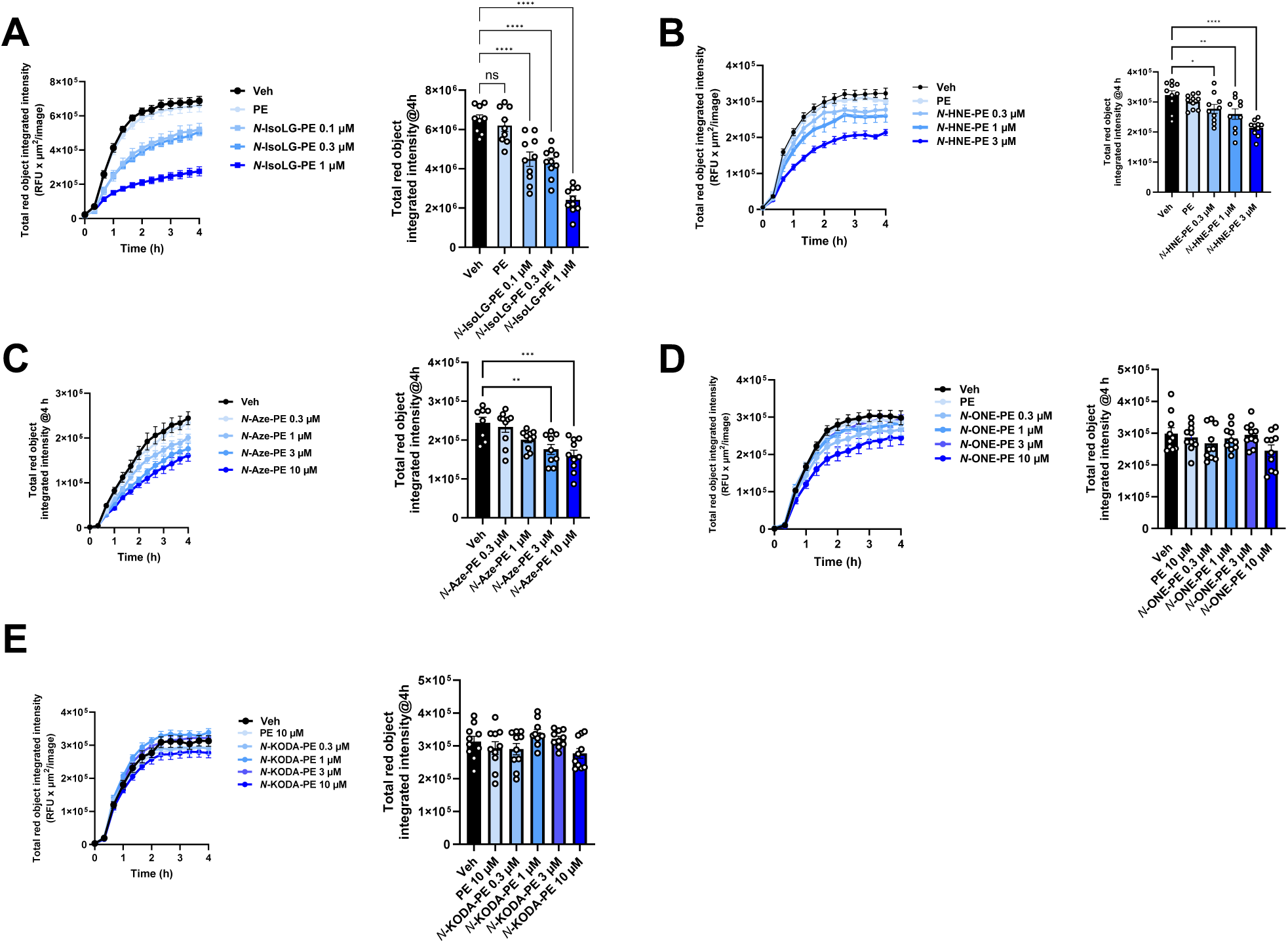
NAMPs inhibit efferocytosis by macrophages. Bone marrow-derived macrophages (BMDMs) were pretreated with the indicated concentrations of individual NAMP species for 6 h prior to addition of CypHer-labeled apoptotic Jurkat cells. Efferocytosis of labeled apoptotic cells was monitored in real time using the Incucyte S3 Live-Cell Analysis System, and red fluorescence was quantified as total integrated intensity. For each NAMP, the left panel shows efferocytosis kinetics and the right panel shows total integrated red fluorescence at the 4-hour endpoint. Data are presented as mean ± SEM. (A) *N*-IsoLG-PE, 1-way ANOVA p<0.0001; (B) *N*-HNE-PE, 1-way ANOVA p<0.0001; (C) *N-*Aze-PE, 1-way ANOVA p<0.0001; (D) *N*-ONE-PE, 1-way ANOVA p=0.1561; and (E) *N*-KODA-PE, 1-way ANOVA, p=0.0576; Dunnett’s multiple-comparison testing vs vehicle, *p < 0.05, **P < 0.01, ***P < 0.001, ****P < 0.0001.

NAPE-PLD hydrolyzes NAMPs which could potentially degrade their efferocytosis-inhibiting properties. To test this, submaximal concentrations of *N*-IsoLG-PE, *N*-HNE-PE, and *N*-Aze-PE were preincubated for two hours either with recombinant mouse NAPE-PLD (rNAPE-PLD) or with heat-inactivated recombinant NAPE-PLD (ΔNAPE-PLD). Prior to these studies, we confirmed that *N*-Aze-PE was also a robust substrate for NAPE-PLD (**Supplemental Figure 2**), as synthetic *N*-Aze-PE had not been available for testing when we conducted our previous studies^16^. After extraction of treated NAMPs to remove NAPE-PLD, the extracted hydrolysis mixture was used to treat BMDM for six hours prior to addition of apoptotic cells. Preincubation with active NAPE-PLD, but not heat-inactivated NAPE-PLD, eliminated the efferocytosis-inhibiting properties of *N*-IsoLG-PE (**Fig. 3A**), *N*-HNE-PE (**Fig. 3B**), and *N*-Aze-PE (**Fig. 3C**), demonstrating that only intact NAMPs inhibited efferocytosis. To assess the effect of deleting endogenous expression of NAPE-PLD on the potency of NAMPs to inhibit efferocytosis, BMDM from *Napepld^-/-^* (KO) mice and *Napepld^+/+^* (WT) mice were treated with submaximal concentrations of *N*-IsoLG-PE or *N*-Aze-PE. The relative inhibition of efferocytosis induced by these NAMPs (compared to vehicle treated BMDM of the same genotype) was greater for KO BMDM than WT BMDM (**Fig. 4A,B**). These results are consistent with the notion that NAPE-PLD present in macrophages reduces the inhibitory effects of NAMPs by hydrolyzing them.

**Figure 3.**
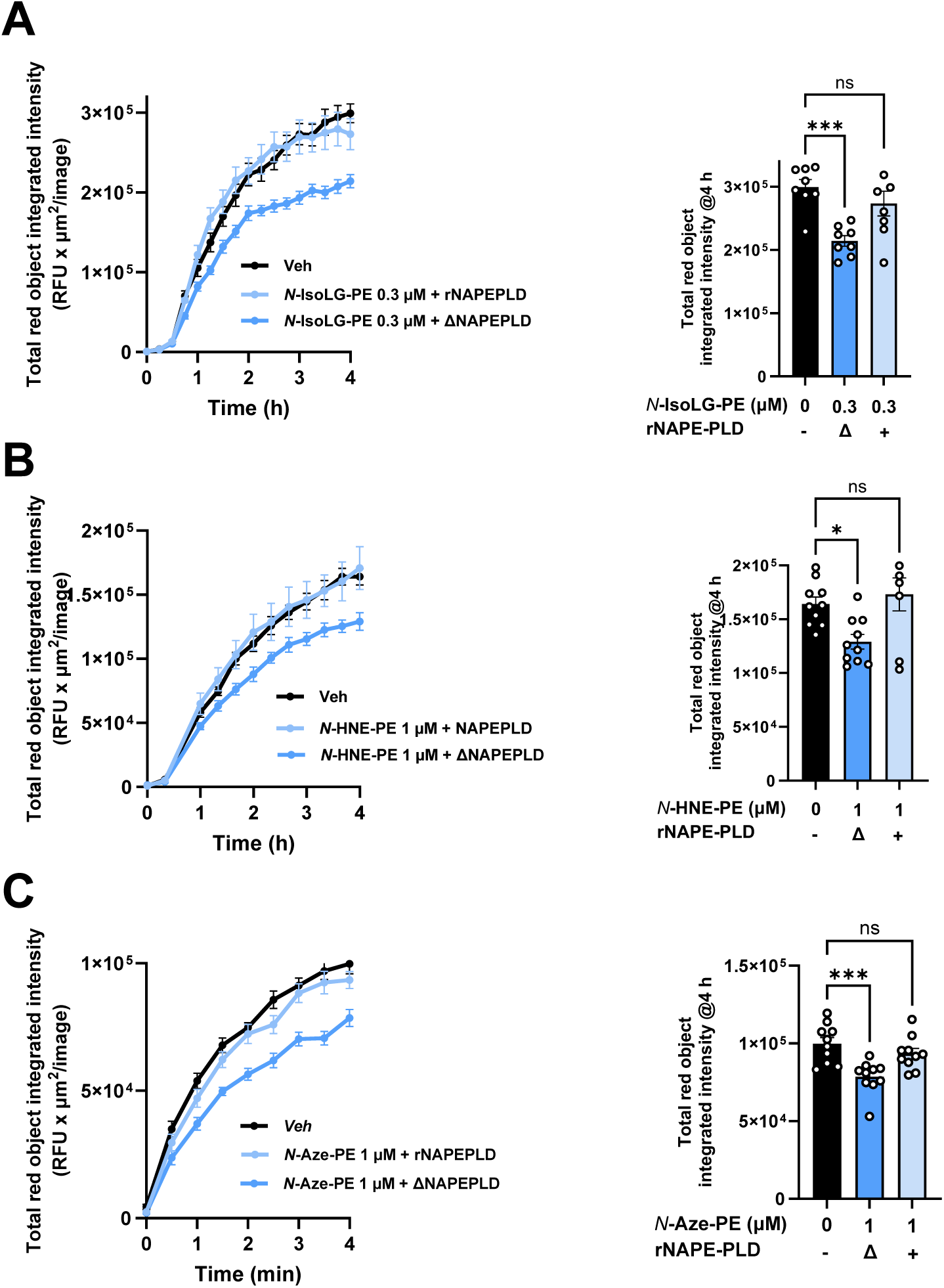
NAPE-PLD hydrolysis abolishes the efferocytosis-inhibiting effects of *N*-aldehyde-modified phosphatidylethanolamines (NAMPs). *N*-IsoLG-PE, *N*-HNE-PE, or *N*-Aze-PE were incubated with active recombinant NAPE-PLD (rNAPE-PLD) or inactive recombinant NAPE-PLD (ΔrNAPE-PLD), followed by lipid extraction to remove enzyme. After treatment with vehicle (0 µM) or NAMP hydrolysis mixture for 6 h, the effect on efferocytosis was assessed by addition of apoptotic Jurkat cells. Left panels: efferocytosis time course. Right panels: total red object integrated red fluorescence at the 4-hour endpoint. (A) 0.3 μM *N*-IsoLG-PE, 1-way ANOVA p=0.0011; (B) 1 μM *N*-HNE-PE, 1-way ANOVA p=0.0073; or (C) 1 μM *N*-Aze-PE, 1-way ANOVA p=0.0008. For each NAMP, Dunnett’s multiple-comparisons test vs vehicle, *p<0.05, ***p<0.001. Data are presented as mean ± SEM.

**Figure 4.**
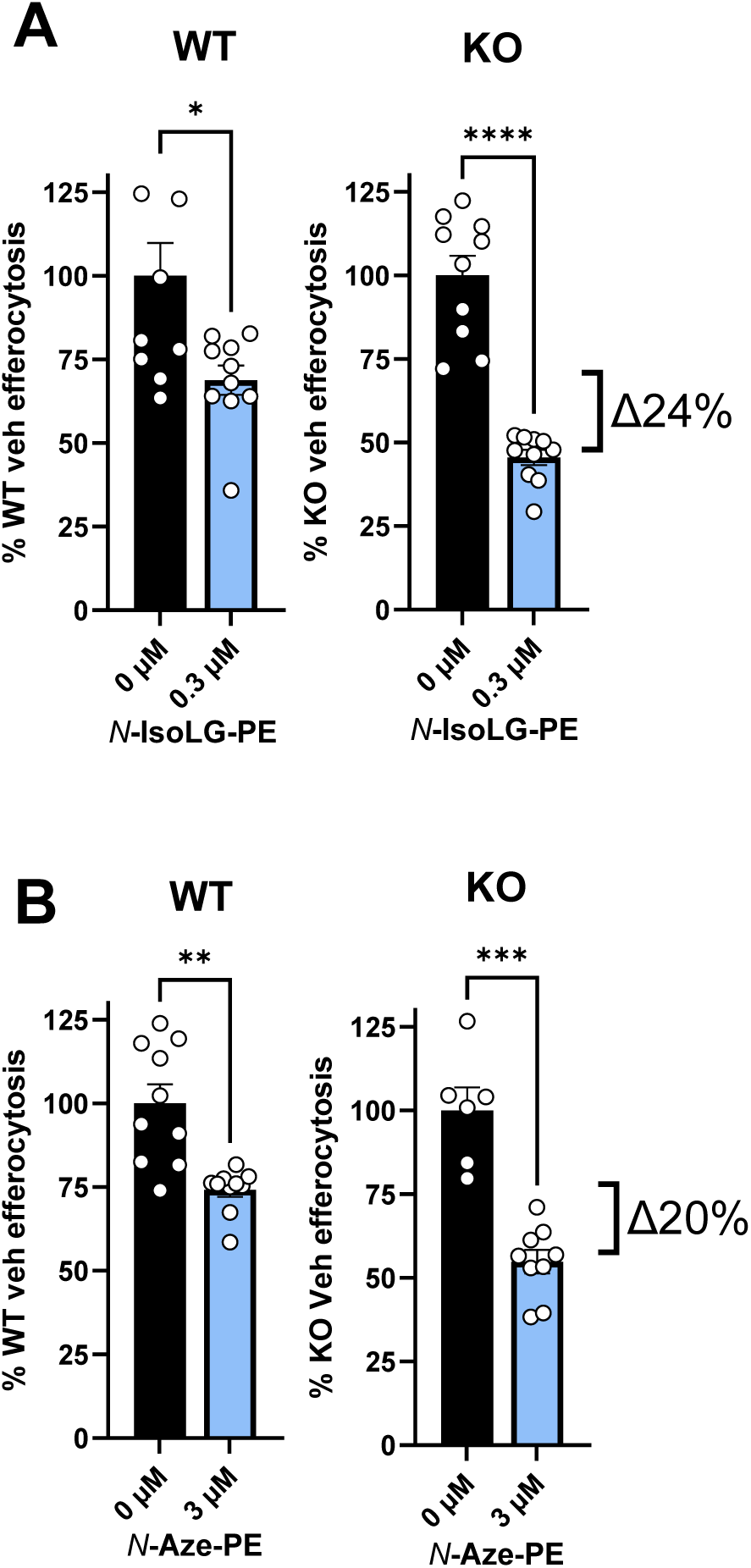
NAPE-PLD deficiency enhances NAMP-mediated inhibition of efferocytosis. *Napepld^+/+^* (WT) and *Napepld^−/−^* (KO) BMDMs were treated with vehicle (0 μM) or (A) 0.3 μM *N*-IsoLG-PE or (B) 3 μM *N*-Aze-PE for 6 h prior to addition of apoptotic Jurkat cells. Efferocytosis is expressed as a percentage of vehicle-treated cells of the same genotype, mean ± SEM, Mann-Whitney test, *p<0.05, **p<0.01, ***p<0.001, ****p<0.0001. Bracket values represent differences in mean %inhibition from vehicle for KO vs WT.

Under atherosclerotic conditions, macrophages might be exposed to elevated levels of NAMPs by other mechanisms in addition to reduced NAPE-PLD expression. For instance, decreased expression of GPX4 is observed in atherosclerotic lesions^29^. GPX4 reduces lipid hydroperoxides to lipid hydroxides and thereby inhibits lipid peroxidation. Thus, modest loss of GPX4 activity promotes lipid peroxidation, while more robust inhibition/deletion of GPX4 triggers ferroptosis (i.e. lipid peroxidation mediated cell death)^30^. To explore the effect of modestly decreased GPX4 activity on NAMP formation and efferocytosis, we first identified 0.3 and 1.0 µM to be sub-lethal concentrations of the GPX4 inhibitor RSL3 (i.e. these concentrations did not cause overt cytotoxicity to BMDM when cytotoxicity was measured by the MTT conversion assay). Even 0.3 µM RSL3 was sufficient to significantly increase cellular levels of NAMPs in BMDM treated with RSL3 for six hours (**Fig. 5A-E**). Treatment of BMDM with 0.3 or 1.0 µM RSL3 also markedly inhibited efferocytosis (**Fig. 5F, G**).

**Figure 5.**
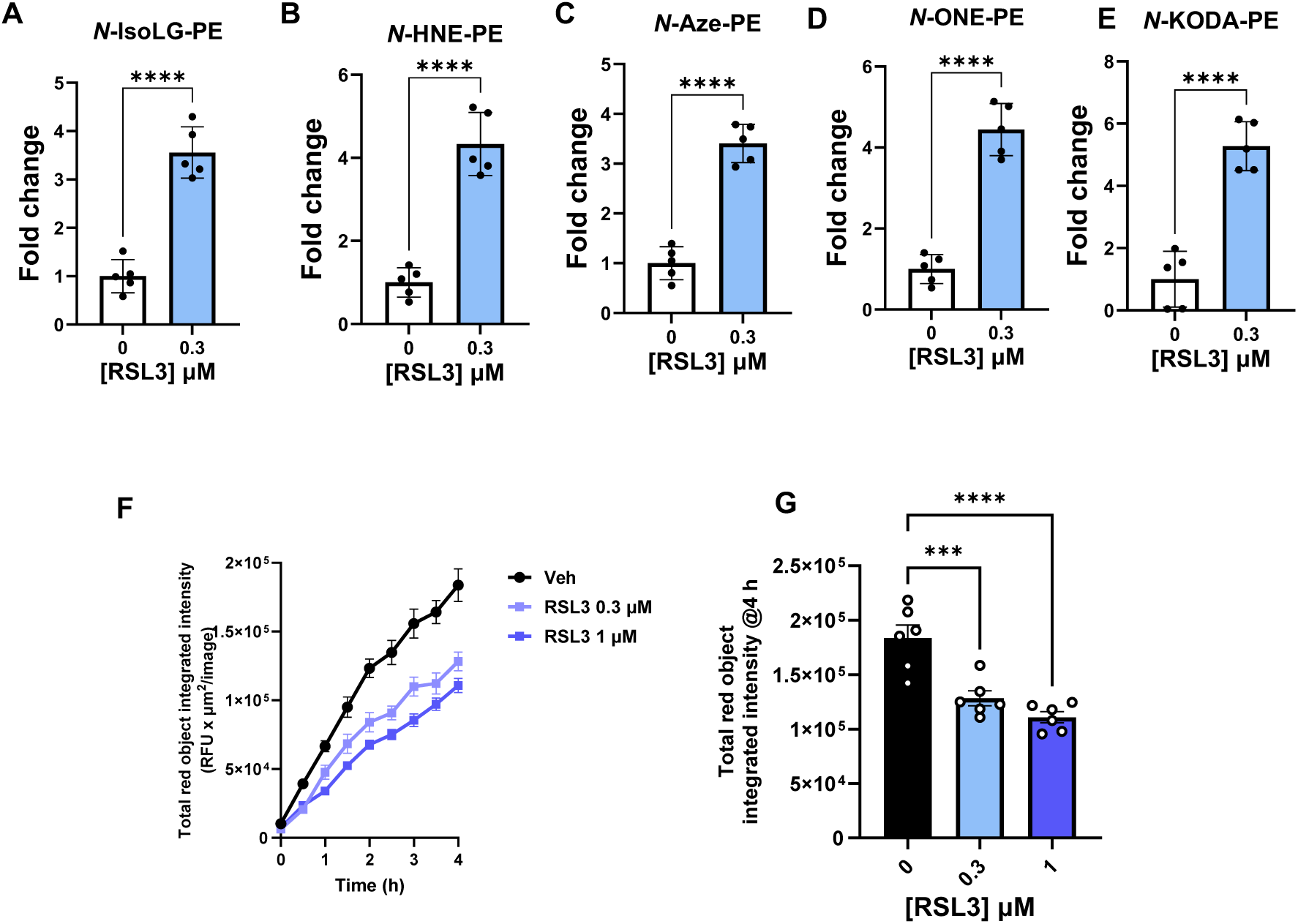
Inhibition of GPX4 by RSL3 increases levels of individual NAMP species in BMDM and inhibits efferocytosis. (A-E) Relative levels of each NAMP after treatment of BMDM with 0.3 µM RSL3 for 6 h was measured by LC/MS. ****p<0.0001, unpaired two-tail t-test. (B) Effects of 6 h pretreatment with 0.3 or 1.0 µM RSL3 on efferocytosis. Left panel: time course; right panel: total red object integrated red fluorescence at the 4-hour endpoint. Mean ± SEM, 1-way ANOVA, p<0.0001; Dunnett’s multiple comparison test vs vehicle, ***p <0.001; ****p<0.0001.

Atherosclerotic conditions could also increase macrophage exposure to NAMPs as the result of oxidation of lipoproteins such as LDL and HDL. Prior studies showed that HDL isolated from individuals with familial hypercholesterolemia have higher levels of *N*-IsoLG-PE^17^ and that in vitro oxidation of HDL by myeloperoxidase produces *N*-IsoLG-PE, along with other NAMPs^14^. To assess whether modification of HDL by reactive lipid aldehydes altered efferocytosis, HDL was pre-treated with varying concentrations of IsoLG, HNE, ONE, and KODA. Because reactions of 9-oxononanoic acid/azeloyl aldehyde acid with PE do not produce high yields of *N*-Aze-PE in the absence of oxidizing reagents, HDL was not pretreated with this aldehyde. Serum-free conditions were used in these assays because serum contains significant amounts of HDL. Treatment of BMDM with 20 µg/mL of unmodified HDL markedly enhanced the ability of BMDM to carry out efferocytosis compared to vehicle treatment (**Fig. 6**). Modification of HDL by IsoLG (**Fig. 6A**) and HNE (**Fig. 6B**), and to a much lesser extent ONE (**Fig. 6C**) and KODA (**Fig. 6D**), resulted in concentration-dependent inhibition of efferocytosis in serum-free conditions, supporting the notion that the oxidative modification of HDL by reactive lipid aldehydes to produce NAMPs could be a driver of impaired efferocytosis.

**Figure 6.**
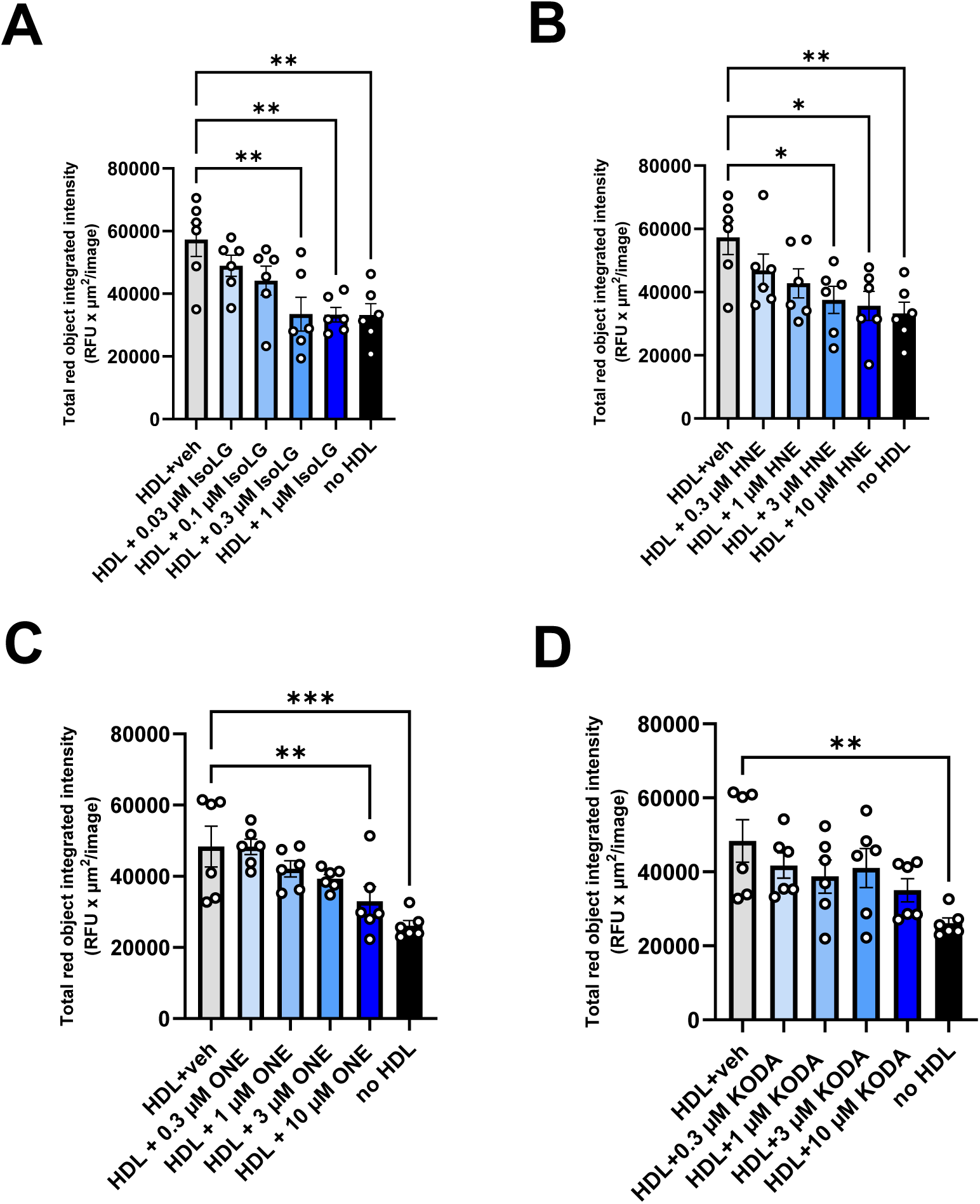
HDL modified by reactive lipids aldehydes inhibits efferocytosis. HDL was incubated with vehicle or varying concentrations of each reactive aldehyde overnight and then the resulting modified HDL incubated with BMDM for 6 h prior to assessing efferocytosis. Data presented as mean ± SEM of total red object integrated intensity at 4-h time point. (A) IsoLG modified HDL, 1-way ANOVA p=0.0009; (B) HNE modified HDL, 1-way ANOVA p=0.0105; (C), ONE modified HDL, 1-way ANOVA p=0.0001; and (D) KODA modified HDL, 1-way ANOVA p=0.0190; for each, Dunnet’s multiple comparisons test vs unmodified HDL *p<0.05, **p<0.01, ***p<0.001.

We recently comprehensively profiled the most abundant NAMP species generated by lipid peroxidation^16^. We therefore quantified these NAMP species in HDL isolated from individuals with FH and control individuals. FH HDL showed significantly increased levels of *N*-HNE-PE, *N*-Aze-PE, *N*-ONE-PE, *N*-IsoLG-PE, and *N*-KODA-PE (**Fig. 7A**). In vitro peroxidation of HDL from control individuals with the radical initiator V70 increased the levels of many of these same NAMPs compared to unoxidized HDL (**Fig. 7B**). Compared to unoxidized HDL, oxidation of HDL by V70 severely reduced its ability to enhance efferocytosis in serum-free conditions (**Fig. 7C**). Performing the V70 oxidation of HDL in the presence of 2HOBA, a highly effective scavenger of reactive lipid aldehydes^31^, partially prevented the inhibitory effects of HDL oxidation, supporting the role of lipid aldehyde adducts including NAMPs in mediating the inhibitory effects of oxidized HDL (**Fig. 7C**). Treating V70-oxidized HDL with recombinant NAPE-PLD (rPLD) but not heat-inactivated recombinant NAPE-PLD (ΔrPLD), also partially reversed the inhibitory effects of oxidized HDL on efferocytosis (**Fig. 7C**), further supporting the notion that NAMPs mediate the efferocytosis-inhibiting effects of oxidized HDL.

**Figure 7.**
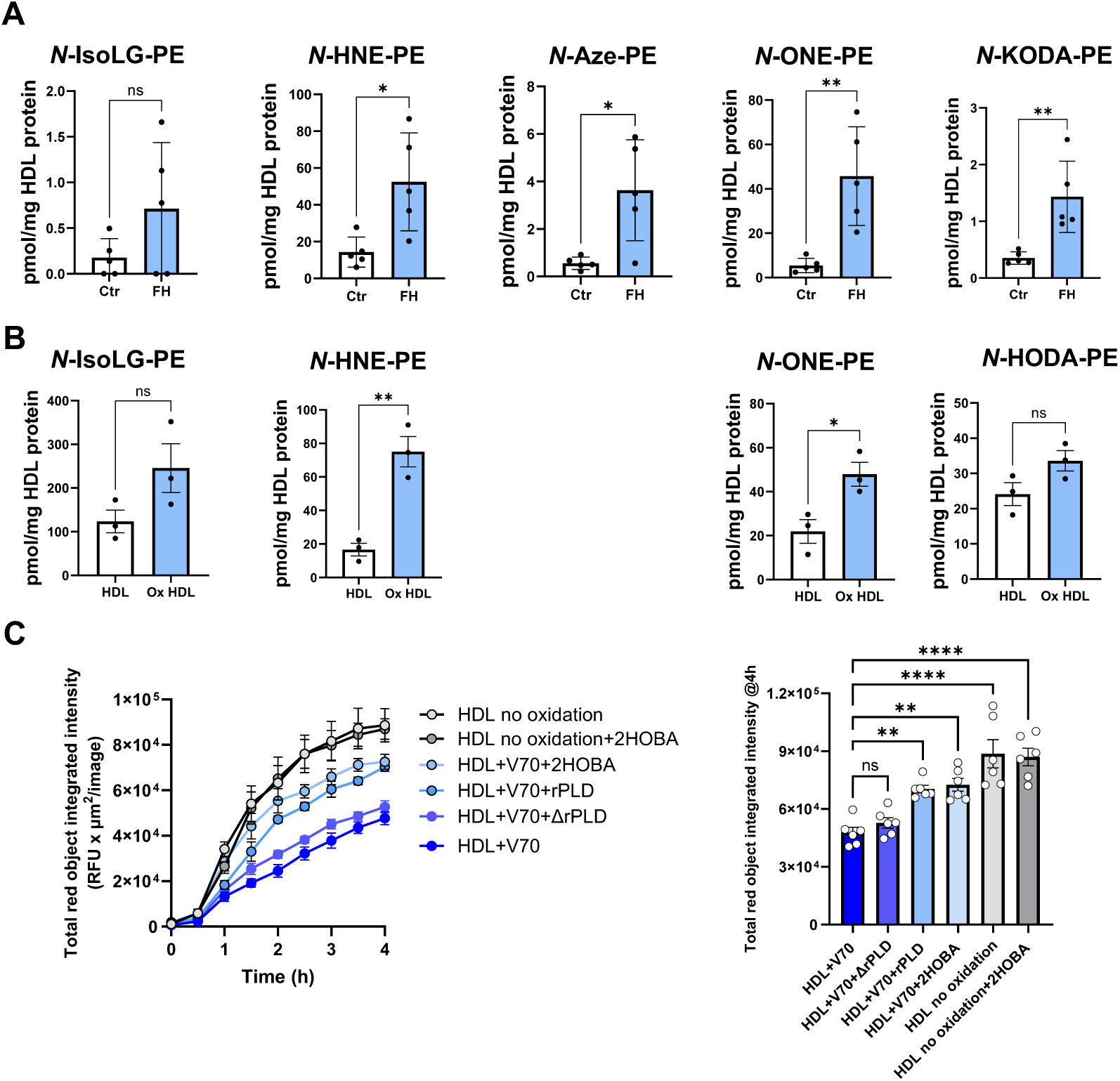
HDL exposed to oxidative environments has higher NAMP levels and inhibits efferocytosis. (A-B) Levels of various NAMP species in isolated HDL were measured by LC/MS/MS after methylamine hydrolysis to remove O-acyl chains. (A) HDL isolated from individuals diagnosed with familial hypercholesterolemia (FH) or control individuals (Ctr). (B) Levels of various NAMP species in HDL incubated with (oxHDL) or without (HDL) the diazo radical initiator V70. Data presented as mean ± SEM, unpaired 2-tailed t-test, *p<0.05, **p<0.001. (C) Effect of oxidized HDL on efferocytosis and effect of various treatments. HDL was peroxidized by diazo initiator V70 in presence or absence of aldehyde scavenger (2HOBA). Additionally, aliquots of V70 peroxidized HDL were subsequently incubated with active recombinant NAPE-PLD (rPLD) or heat inactivated enzyme (ΔrPLD). 20 µg/mL of each HDL preparation was then incubated with BMDM for 6 h before efferocytosis measured by addition of CypHer-labeled apoptotic cells. Left panel: efferocytosis time course. Right panel: total red object integrated red fluorescence at the 4-hour endpoint. Data represents mean ± SEM, 1-way ANOVA, p<0.0001, Dunnett’s multiple comparisons test vs HDL+V70, **p<0.01, ****p<0.0001.

We investigated several potential mechanisms by which NAMPs might inhibit efferocytosis. TNF directly inhibits efferocytosis by macrophages^32^, and various NAMPs have previously been shown to induce pro-inflammatory responses including the secretion of TNF via activation of NFkB^17^. However, pre-treatment of BMDM with the IκK inhibitor BMS-345541 did not block the inhibition of efferocytosis by *N*-IsoLG-PE (**Fig. 8A**) or *N*-Aze-PE (**Fig. 8B**), suggesting that NAMP inhibition of efferocytosis does not depend on NFκB activation. Excessive ROS formation markedly inhibits efferocytosis and prior studies showed that *N*-IsoLG-PE induces increased ROS formation by isolated kidney mitochondria^19^. We therefore assessed the effect of *N*-IsoLG-PE and *N*-Aze-PE on total cellular superoxide levels in BMDM. While *N*-IsoLG-PE significantly increased superoxide levels compared to vehicle treated BMDM, *N*-Aze-PE significantly decreased superoxide formation (**Fig. 8C**). Therefore, either NAMP inhibition of efferocytosis does not depend on the modulation of superoxide levels or the two NAMPs significantly differ in the mechanisms by which they inhibit efferocytosis. Sustained continuous efferocytosis requires efficient cholesterol efflux because the uptake of apoptotic cells strikingly increases the cholesterol content of macrophages^33, 34^. To test if NAMPs hindered the ability of BMDM to efficiently efflux cholesterol, BMDM were loaded with BODIPY-cholesterol and then treated with media containing 10% serum and vehicle, *N*-IsoLG-PE or *N*-Aze-PE for 6 h and the extent of BODIPY-cholesterol transfer from BMDM to the media was measured by fluorescence. Both *N*-IsoLG-PE (**Fig. 8D**) and *N*-Aze-PE (**Fig. 8E**) markedly inhibited cholesterol efflux, supporting inhibition of cholesterol efflux as one mechanism by which NAMPs inhibit efferocytosis.

**Figure 8.**
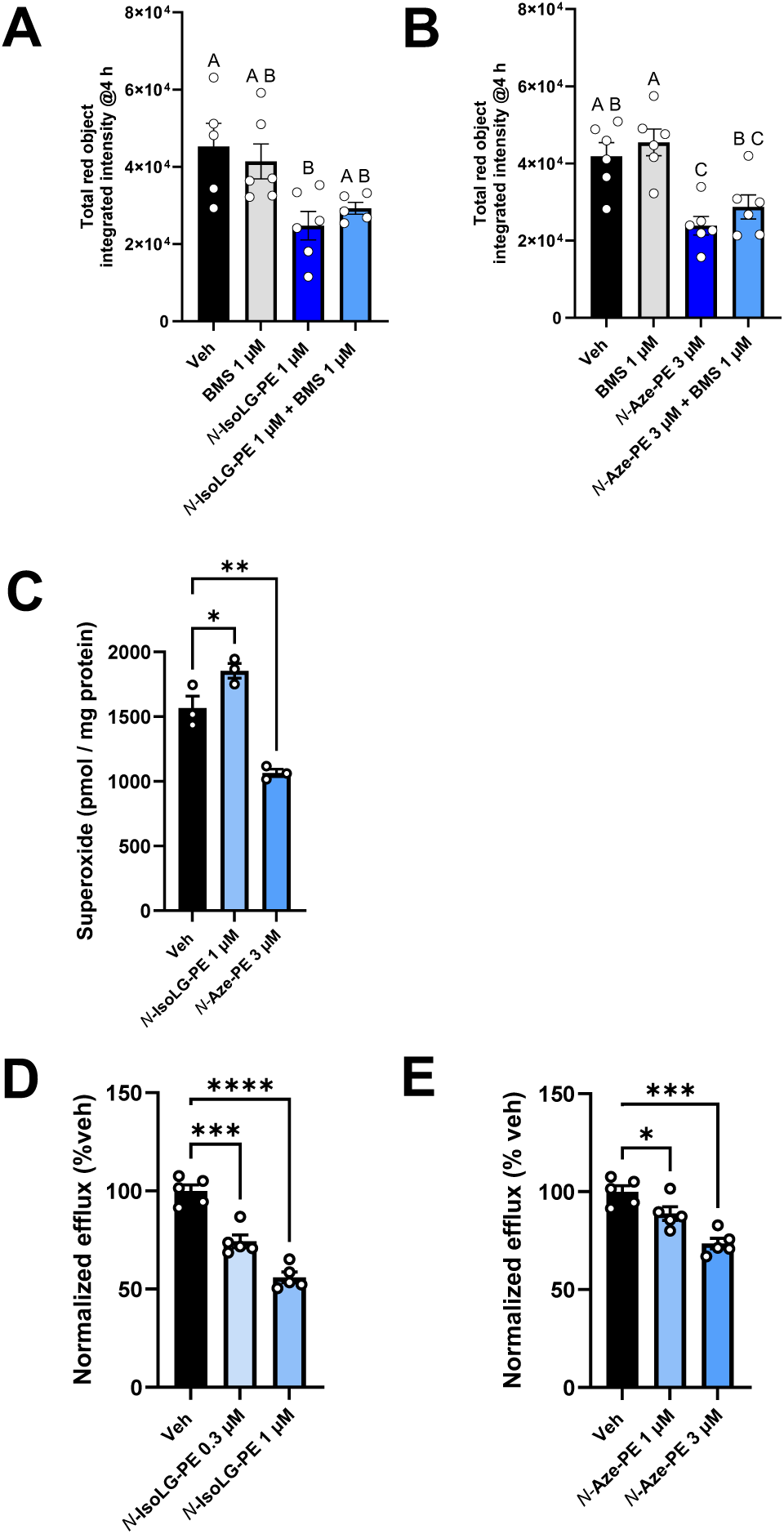
Assessment of potential mechanisms for NAMP inhibition of efferocytosis. (A-B) BMDM were pretreated with IKK inhibitor BMS-345541 for 1 h prior to treatment for 6 h with either (A) 1 µM *N*-IsoLG-PE, 1-way ANOVA p= 0.0093 or (B) 3 µM *N*-Aze-PE, 1-way ANOVA p= 0.0020. Sidak’s multiple comparisons test, treatments with same letter are not significantly different from each other. (C) Following 6 h treatment with vehicle, *N*-IsoLG-PE, or *N*-Aze-PE, cells were washed and superoxide present in cell measured by electron spin resonance with TMH oxidation as probe. 1-way ANOVA, p=0.0004, Dunnett’s multiple comparisons test vs veh, *p<0.05, **p<0.001. (D-E) Effect of incubating media containing 10% serum and either vehicle (veh) or NAMPs for 6 h on cholesterol efflux to media from BODIPY-cholesterol loaded BMDM. Efflux was normalized to BMDM treated with media with vehicle only. (D) *N*-IsoLG-PE, 1-way ANOVA p<0.0001 and (E) *N*-Aze-PE, 1-way ANOVA p=0.0002. Dunnett’s multiple comparisons test vs veh, *p<0.05, ***p<0.001, ****p<0.0001. All data are presented as mean ± SEM.

## DISCUSSION

Both enhanced lipid peroxidation^35–38^ and impaired efferocytosis^39–41^ are consistent features of chronic inflammatory diseases including atherosclerosis, diabetic skin wounds, systemic lupus erythematosus, rheumatoid arthritis, and inflammatory bowel diseases. Our findings that conditions that increase lipid peroxidation (HDL oxidation, familial hypercholesterolemia, and GPX4 inhibition) significantly increased NAMP levels and impaired efferocytosis and that exposure to NAMPs was sufficient to significantly inhibit efferocytosis, suggest that the two are causally related, with lipid peroxidation inhibiting efferocytosis at least in part via NAMP formation. That NAMPs are indeed elevated *in vivo* under relevant conditions is shown by our finding of elevated NAMP levels in HDL isolated from individuals with familial hypercholesterolemia. Although rigorous proof that NAMP formation impairs efferocytosis *in vivo* will require additional studies showing that interventions that reduce NAMP levels (e.g. aldehyde scavengers, rNAPE-PLD, or NAPE-PLD activators) consistently restore efficient efferocytosis, it is worth noting that treatment of atheroprone *Ldlr^-/-^*mice with the aldehyde scavenger 2HOBA has already been shown to improve *in vivo* measures of efferocytosis^42^, although the effect of 2HOBA treatment on NAMPs levels was not determined in that study.

While lipid peroxidation generates many species of peroxidized lipids, our finding that pretreatment of oxidized HDL with NAPE-PLD, which specifically degrades NAMPs, markedly diminishes the inhibitory effect of oxidized HDL on efferocytosis implicates NAMPs as being significant contributors to this inhibition. Prior studies showed that NAPE-PLD degrades several abundant NAMPs such as *N*-IsoLG-PE, *N*-HNE-PE, and *N-*KODA-PE^16, 43^ and this current study demonstrates that NAPE-PLD also degrades *N*-Aze-PE. Our finding that pretreatment with recombinant NAPE-PLD prevented, while genetic deletion of NAPE-PLD potentiated, the inhibitory effects of various NAMPs on efferocytosis is consistent with the degradation of NAMPs being a biologically important function of NAPE-PLD and that the reduced capacity for efferocytosis previously observed with NAPE-PLD deletion or inhibition^43^ arises at least in part from the ensuing accumulation of NAMPs.

In a variety of chronic inflammatory diseases where impaired efferocytosis is observed, decreased expression and/or activity of GPX4 has also been reported including atherosclerotic lesions^29^, inflammatory bowel disease^44^, rheumatoid arthritis^45^, systemic lupus erythematosus^46^ and diabetic kidney disease^47^. Our finding that even modest inhibition of GPX4 (i.e. below that needed to induce ferroptosis) increased formation of NAMPs within macrophages and markedly inhibited efferocytosis, supports a causative relationship between NAMP elevations and impaired efferocytosis when GPX4 is reduced. These findings might also explain the observation that ferroptotic cells are cleared less efficiently than apoptotic cells^48^, since apoptotic cells with intact GPX4 would presumably have much lower levels of NAMPs.

Additional studies are required to fully elucidate the mechanisms by which NAMPs inhibit efferocytosis. This study examined the potential contribution of three key pathways known to regulate efferocytosis: NFκB activation, ROS production, and cholesterol efflux. Activation of NFκB plays an important role in polarization of macrophages towards a pro-inflammatory phenotype (M1-like), which tend to have reduced efferocytosis capacity^49, 50^. For instance, treatment of macrophages with LPS or TNF induces NFκB and inhibits efferocytosis^51^. Prior studies found that various NAMPs induce pro-inflammatory responses with *N*-IsoLG-PE inducing TNF via the receptor for advanced glycation end-products (RAGE)^17^. However, blocking NFκB activation by treating with the IκK inhibitor BMS-345541 failed to protect against NAMP-induced inhibition of efferocytosis, suggesting that NFκB activation is not required. Elevated ROS production inhibits efferocytosis ^52, 53^. Superoxide production by NADPH oxidase has also been reported to compete with phagolysosome acidification^54^, a key step in efferocytosis. However, efferocytosis has also been reported to activate NADPH oxidase, and macrophages without NADPH oxidase show delayed maturation of efferosomes^55^. Thus, how changes in ROS production will affect efferocytosis may be difficult to predict. In any case, the effect of various inhibitory NAMPs on superoxide production does not appear to be consistent, with *N*-IsoLG-PE elevating superoxide levels and *N*-Aze-PE decreasing them. For this reason, we consider increased ROS production unlikely to be the unified mechanism by which NAMPs inhibit efferocytosis. Changes in macrophages cholesterol efflux capacity consistently correlate with changes in their efferocytosis capacity^33, 34, 56–61^. Macrophage uptake of apoptotic cells markedly increases their cholesterol content, so that in the absence of efficient cholesterol efflux, the resulting cholesterol overload hampers phagocytosis and triggers apoptosis^33^. ABCA1plays a key role in cholesterol efflux as well in the recognition and clearance of apoptotic cells through regulation of both “find-me” and “eat-me” signaling pathways^61^. We found that BMDMs treated with *N*-Aze-PE or *N*-IsoLG-PE cannot efficiently efflux cholesterol, suggesting this in an important mechanism contributing to NAMP-induced inhibition of efferocytosis. Precisely how NAMPs inhibit cholesterol efflux remains to be determined.

In summary, these studies demonstrate that processes highly relevant to chronic inflammation markedly increase NAMP levels and that multiple NAMP species inhibit efferocytosis, so that elevated levels of NAMPs can plausibly contribute to the impaired efferocytosis observed in various chronic inflammatory conditions. If so, then preventing the formation of NAMPs or enhancing the rate of NAMP degradation may be important new approaches to rescuing this impaired efferocytosis.

## Author contributions

All authors reviewed the manuscript. R.F. assisted with conceptualization, performed the majority of the experiments, analyzed data, developed the LC/MS methodology, prepared figures and assisted in the writing and editing of the original draft of the manuscript. A.B., E.A.W., Z.M., and AMAO assisted with experiments. A.L.G. assisted with experiments and in the development of the *in vitro* efferocytosis method. KAT synthesized *N*-Aze-PE and edited the manuscript. S.D. performed the superoxide measurements and edited the manuscript. M.F.L. assisted in funding acquisition, recruited FH and control patients, supervised blood collection, and edited the manuscript. K.C. Vickers isolated HDL from patient plasma and edited the manuscript. A.C.D assisted in conceptualization of studies, supervised the development and execution of in vitro efferocytosis studies, provided reagents, and assisted in the preparation of figures and editing of the manuscript. S.S.D. conceptualized the studies, supervised and administered the overall project, assisted in funding acquisition, assisted in data analysis and development of methodology, assisted in preparation of figures, and wrote the original draft of the manuscript.

## Acknowledgments

Funding was provided in part by a program project grant from the National Heart, Blood, and Lung Institute P01HL116263-06A1 and from institutional funds provided by Vanderbilt University.

## Ethics Approval

All animal procedures were approved by the Vanderbilt University Institutional Animal Care and Use Committee and conducted in accordance with the Guide for Care and Use of Laboratory Animals. The collection of blood from patients with familial hypercholesterolemia and control individuals was approved by the Vanderbilt University Medical Center Institutional Review Board. All participants provided written consent prior to enrollment.

## References

1. Zhang B, Zou Y, Yuan Z, Jiang K, Zhang Z, Chen S, Zhou X, Wu Q, Zhang X. Efferocytosis: the resolution of inflammation in cardiovascular and cerebrovascular disease. Front Immunol. 2024;15:1485222. Epub 20241126. doi: 10.3389/fimmu.2024.1485222. PubMed PMID: 39660125; PMCID: PMC11628373.

2. Doran AC. Inflammation Resolution: Implications for Atherosclerosis. Circ Res. 2022;130(1):130–48. Epub 20220107. doi: 10.1161/circresaha.121.319822. PubMed PMID: 34995137; PMCID: PMC8842990.

3. Ge Y, Huang M, Yao YM. Efferocytosis and Its Role in Inflammatory Disorders. Front Cell Dev Biol. 2022;10:839248. Epub 20220225. doi: 10.3389/fcell.2022.839248. PubMed PMID: 35281078; PMCID: PMC8913510.

4. Wang L, Li H, Tang Y, Yao P. Potential Mechanisms and Effects of Efferocytosis in Atherosclerosis. Front Endocrinol (Lausanne). 2020;11:585285. Epub 20210201. doi: 10.3389/fendo.2020.585285. PubMed PMID: 33597922; PMCID: PMC7883484.

5. Analysis of the 17β-estradiol (17βE2)-, and coumestrol-regulated transcriptomes in MCF-7 cells. In: Signaling Pathways P, editor.: Signaling Pathways Project; 2009.

6. Akyol O, Chiang HH, Burns AR, Yang CY, Woodside DG, Sawamura T, Sánchez-Quesada JL, Gotto AM, Chen CH. LDL atherogenicity determined by size, density, oxidation, apolipoprotein(a), and electronegativity: an updated review. Front Cardiovasc Med. 2025;12:1649759. Epub 20251024. doi: 10.3389/fcvm.2025.1649759. PubMed PMID: 41210345; PMCID: PMC12592080.

7. Tsimikas S, Witztum JL. Oxidized phospholipids in cardiovascular disease. Nat Rev Cardiol. 2024;21(3):170–91. Epub 20231017. doi: 10.1038/s41569-023-00937-4. PubMed PMID: 37848630.

8. Esterbauer H, Schaur RJ, Zollner H. Chemistry and biochemistry of 4-hydroxynonenal, malonaldehyde and related aldehydes. Free Radic Biol Med. 1991;11(1):81–128. doi: 10.1016/0891-5849(91)90192-6. PubMed PMID: 1937131.

9. Fritz KS, Petersen DR. An overview of the chemistry and biology of reactive aldehydes. Free Radic Biol Med. 2013;59:85–91. Epub 20120628. doi: 10.1016/j.freeradbiomed.2012.06.025. PubMed PMID: 22750507; PMCID: PMC3540155.

10. Lee SH, Blair IA. Characterization of 4-oxo-2-nonenal as a novel product of lipid peroxidation. Chem Res Toxicol. 2000;13(8):698–702. doi: 10.1021/tx000101a. PubMed PMID: 10956056.

11. Podrez EA, Poliakov E, Shen Z, Zhang R, Deng Y, Sun M, Finton PJ, Shan L, Gugiu B, Fox PL, Hoff HF, Salomon RG, Hazen SL. Identification of a novel family of oxidized phospholipids that serve as ligands for the macrophage scavenger receptor CD36. J Biol Chem. 2002;277(41):38503–16. Epub 20020708. doi: 10.1074/jbc.M203318200. PubMed PMID: 12105195.

12. Schneider C, Tallman KA, Porter NA, Brash AR. Two distinct pathways of formation of 4-hydroxynonenal. Mechanisms of nonenzymatic transformation of the 9- and 13-hydroperoxides of linoleic acid to 4-hydroxyalkenals. J Biol Chem. 2001;276(24):20831–8. Epub 20010319. doi: 10.1074/jbc.M101821200. PubMed PMID: 11259420.

13. Brame CJ, Salomon RG, Morrow JD, Roberts LJ, 2nd. Identification of extremely reactive gamma-ketoaldehydes (isolevuglandins) as products of the isoprostane pathway and characterization of their lysyl protein adducts. J Biol Chem. 1999;274(19):13139–46. doi: 10.1074/jbc.274.19.13139. PubMed PMID: 10224068.

14. Guo L, Chen Z, Amarnath V, Davies SS. Identification of novel bioactive aldehyde-modified phosphatidylethanolamines formed by lipid peroxidation. Free Radic Biol Med. 2012;53(6):1226–38. Epub 20120804. doi: 10.1016/j.freeradbiomed.2012.07.077. PubMed PMID: 22898174; PMCID: PMC3461964.

15. Davies SS, Guo L. Lipid peroxidation generates biologically active phospholipids including oxidatively N-modified phospholipids. Chem Phys Lipids. 2014;181:1–33. Epub 20140402. doi: 10.1016/j.chemphyslip.2014.03.002. PubMed PMID: 24704586; PMCID: PMC4075969.

16. Fadaei R, Bernstein AC, Jenkins AN, Pickens AG, Zarrow JE, Alli-Oluwafuyi AM, Tallman KA, Davies SS. N-aldehyde-modified phosphatidylethanolamines generated by lipid peroxidation are robust substrates of N-acyl phosphatidylethanolamine phospholipase D. J Lipid Res. 2025;66(7):100831. Epub 20250521. doi: 10.1016/j.jlr.2025.100831. PubMed PMID: 40409473; PMCID: PMC12214272.

17. Guo L, Chen Z, Amarnath V, Yancey PG, Van Lenten BJ, Savage JR, Fazio S, Linton MF, Davies SS. Isolevuglandin-type lipid aldehydes induce the inflammatory response of macrophages by modifying phosphatidylethanolamines and activating the receptor for advanced glycation endproducts. Antioxid Redox Signal. 2015;22(18):1633–45. Epub 20150318. doi: 10.1089/ars.2014.6078. PubMed PMID: 25751734; PMCID: PMC4485367.

18. Guo L, Chen Z, Cox BE, Amarnath V, Epand RF, Epand RM, Davies SS. Phosphatidylethanolamines modified by γ-ketoaldehyde (γKA) induce endoplasmic reticulum stress and endothelial activation. J Biol Chem. 2011;286(20):18170–80. Epub 20110325. doi: 10.1074/jbc.M110.213470. PubMed PMID: 21454544; PMCID: PMC3093889.

19. Mayorov V, Uchakin P, Amarnath V, Panov AV, Bridges CC, Uzhachenko R, Zackert B, Moore CS, Davies S, Dikalova A, Dikalov S. Targeting of reactive isolevuglandins in mitochondrial dysfunction and inflammation. Redox Biol. 2019;26:101300. Epub 20190814. doi: 10.1016/j.redox.2019.101300. PubMed PMID: 31437812; PMCID: PMC6831880.

20. Rinne P, Guillamat-Prats R, Rami M, Bindila L, Ring L, Lyytikäinen LP, Raitoharju E, Oksala N, Lehtimäki T, Weber C, van der Vorst EPC, Steffens S. Palmitoylethanolamide Promotes a Proresolving Macrophage Phenotype and Attenuates Atherosclerotic Plaque Formation. Arterioscler Thromb Vasc Biol. 2018;38(11):2562–75. doi: 10.1161/atvbaha.118.311185. PubMed PMID: 30354245.

21. Zarrow JE, Alli-Oluwafuyi AM, Youwakim CM, Kim K, Jenkins AN, Suero IC, Jones MR, Mashhadi Z, Mackie K, Waterson AG, Doran AC, Sulikowski GA, Davies SS. Small Molecule Activation of NAPE-PLD Enhances Efferocytosis by Macrophages. ACS Chem Biol. 2023;18(8):1891–904. Epub 20230802. doi: 10.1021/acschembio.3c00401. PubMed PMID: 37531659; PMCID: PMC10443532.

22. Okamoto Y, Morishita J, Tsuboi K, Tonai T, Ueda N. Molecular characterization of a phospholipase D generating anandamide and its congeners. J Biol Chem. 2004;279(7):5298–305. Epub 20031121. doi: 10.1074/jbc.M306642200. PubMed PMID: 14634025.

23. Hussain Z, Uyama T, Tsuboi K, Ueda N. Mammalian enzymes responsible for the biosynthesis of N-acylethanolamines. Biochim Biophys Acta Mol Cell Biol Lipids. 2017;1862(12):1546–61. Epub 20170824. doi: 10.1016/j.bbalip.2017.08.006. PubMed PMID: 28843504.

24. Liu J, Wang L, Harvey-White J, Huang BX, Kim H-Y, Luquet S, Palmiter RD, Krystal G, Rai R, Mahadevan A, Razdan RK, Kunos G. Multiple pathways involved in the biosynthesis of anandamide. Neuropharmacology. 2008;54(1):1–7. doi: 10.1016/j.neuropharm.2007.05.020.

25. Leishman E, Mackie K, Luquet S, Bradshaw HB. Lipidomics profile of a NAPE-PLD KO mouse provides evidence of a broader role of this enzyme in lipid metabolism in the brain. Biochimica et Biophysica Acta (BBA) - Molecular and Cell Biology of Lipids. 2016;1861(6):491–500. doi: 10.1016/j.bbalip.2016.03.003.

26. Dikalov SI, Kirilyuk IA, Voinov M, Grigor’ev IA. EPR detection of cellular and mitochondrial superoxide using cyclic hydroxylamines. Free Radic Res. 2011;45(4):417–30. Epub 20101203. doi: 10.3109/10715762.2010.540242. PubMed PMID: 21128732; PMCID: PMC4210377.

27. Dikalov SI, Polienko YF, Kirilyuk I. Electron Paramagnetic Resonance Measurements of Reactive Oxygen Species by Cyclic Hydroxylamine Spin Probes. Antioxid Redox Signal. 2018;28(15):1433–43. Epub 20171117. doi: 10.1089/ars.2017.7396. PubMed PMID: 29037084; PMCID: PMC5910043.

28. Sankaranarayanan S, Kellner-Weibel G, de la Llera-Moya M, Phillips MC, Asztalos BF, Bittman R, Rothblat GH. A sensitive assay for ABCA1-mediated cholesterol efflux using BODIPY-cholesterol. J Lipid Res. 2011;52(12):2332–40. Epub 20110927. doi: 10.1194/jlr.D018051. PubMed PMID: 21957199; PMCID: PMC3220299.

29. Liu Z, Cheng S, Zheng X, Wang X, Lu W, Wang X, Pan L, Shan Y, Qiu C. Paclitaxel Attenuates Atherosclerosis by Suppressing Macrophage Ferroptosis and Improving Lipid Metabolism via the Sirt1/Nrf2/GPX4 Pathway. Faseb j. 2025;39(15):e70917. doi: 10.1096/fj.202501047RR. PubMed PMID: 40779351.

30. Ursini F, Maiorino M. Lipid peroxidation and ferroptosis: The role of GSH and GPx4. Free Radic Biol Med. 2020;152:175–85. Epub 20200309. doi: 10.1016/j.freeradbiomed.2020.02.027. PubMed PMID: 32165281.

31. May-Zhang LS, Kirabo A, Huang J, Linton MF, Davies SS, Murray KT. Scavenging Reactive Lipids to Prevent Oxidative Injury. Annu Rev Pharmacol Toxicol. 2021;61:291–308. Epub 20200930. doi: 10.1146/annurev-pharmtox-031620-035348. PubMed PMID: 32997599.

32. Kojima Y, Volkmer JP, McKenna K, Civelek M, Lusis AJ, Miller CL, Direnzo D, Nanda V, Ye J, Connolly AJ, Schadt EE, Quertermous T, Betancur P, Maegdefessel L, Matic LP, Hedin U, Weissman IL, Leeper NJ. CD47-blocking antibodies restore phagocytosis and prevent atherosclerosis. Nature. 2016;536(7614):86–90. Epub 20160720. doi: 10.1038/nature18935. PubMed PMID: 27437576; PMCID: PMC4980260.

33. Yvan-Charvet L, Pagler TA, Seimon TA, Thorp E, Welch CL, Witztum JL, Tabas I, Tall AR. ABCA1 and ABCG1 protect against oxidative stress-induced macrophage apoptosis during efferocytosis. Circ Res. 2010;106(12):1861–9. Epub 20100429. doi: 10.1161/circresaha.110.217281. PubMed PMID: 20431058; PMCID: PMC2995809.

34. Anandan V, Thulaseedharan T, Suresh Kumar A, Chandran Latha K, Revikumar A, Mullasari A, Kartha CC, Jaleel A, Ramachandran S. Cyclophilin A Impairs Efferocytosis and Accelerates Atherosclerosis by Overexpressing CD 47 and Down-Regulating Calreticulin. Cells. 2021;10(12). Epub 20211220. doi: 10.3390/cells10123598. PubMed PMID: 34944106; PMCID: PMC8700718.

35. Gianazza E, Brioschi M, Martinez Fernandez A, Casalnuovo F, Altomare A, Aldini G, Banfi C. Lipid Peroxidation in Atherosclerotic Cardiovascular Diseases. Antioxid Redox Signal. 2021;34(1):49–98. Epub 20200827. doi: 10.1089/ars.2019.7955. PubMed PMID: 32640910.

36. Feng J, Wang J, Wang Y, Huang X, Shao T, Deng X, Cao Y, Zhou M, Zhao C. Oxidative Stress and Lipid Peroxidation: Prospective Associations Between Ferroptosis and Delayed Wound Healing in Diabetic Ulcers. Front Cell Dev Biol. 2022;10:898657. Epub 20220708. doi: 10.3389/fcell.2022.898657. PubMed PMID: 35874833; PMCID: PMC9304626.

37. Leitinger N. The role of phospholipid oxidation products in inflammatory and autoimmune diseases: evidence from animal models and in humans. Subcell Biochem. 2008;49:325–50. doi: 10.1007/978-1-4020-8830-8_12. PubMed PMID: 18751917.

38. Cracowski JL, Bonaz B, Bessard G, Bessard J, Anglade C, Fournet J. Increased urinary F2-isoprostanes in patients with Crohn’s disease. Am J Gastroenterol. 2002;97(1):99–103. doi: 10.1111/j.1572-0241.2002.05427.x. PubMed PMID: 11808977.

39. Adkar SS, Leeper NJ. Efferocytosis in atherosclerosis. Nat Rev Cardiol. 2024;21(11):762–79. Epub 20240515. doi: 10.1038/s41569-024-01037-7. PubMed PMID: 38750215.

40. Sun Y, Guo H, Bai Y, Chen J, Li Y. Roles of efferocytosis in wound repair: Process, cells, and signals. Genes Dis. 2026;13(3):101937. Epub 20251113. doi: 10.1016/j.gendis.2025.101937. PubMed PMID: 41630951; PMCID: PMC12860986.

41. He Z, Chen N, Zhang Y, Du H, Jie L. Impaired Macrophage Efferocytosis: Shared Mechanisms and Therapeutic Implications in Immune-Mediated Inflammatory Diseases. J Inflamm Res. 2026;19:582297. Epub 20260408. doi: 10.2147/jir.S582297. PubMed PMID: 41978837; PMCID: PMC13070338.

42. Tao H, Huang J, Yancey PG, Yermalitsky V, Blakemore JL, Zhang Y, Ding L, Zagol-Ikapitte I, Ye F, Amarnath V, Boutaud O, Oates JA, Roberts LJ, 2nd, Davies SS, Linton MF. Scavenging of reactive dicarbonyls with 2-hydroxybenzylamine reduces atherosclerosis in hypercholesterolemic Ldlr(-/-) mice. Nat Commun. 2020;11(1):4084. Epub 20200814. doi: 10.1038/s41467-020-17915-w. PubMed PMID: 32796843; PMCID: PMC7429830.

43. Guo L, Gragg SD, Chen Z, Zhang Y, Amarnath V, Davies SS. Isolevuglandin-modified phosphatidylethanolamine is metabolized by NAPE-hydrolyzing phospholipase D. J Lipid Res. 2013;54(11):3151–7. Epub 20130909. doi: 10.1194/jlr.M042556. PubMed PMID: 24018423; PMCID: PMC3793619.

44. Mayr L, Grabherr F, Schwärzler J, Reitmeier I, Sommer F, Gehmacher T, Niederreiter L, He G-W, Ruder B, Kunz KTR, Tymoszuk P, Hilbe R, Haschka D, Feistritzer C, Gerner RR, Enrich B, Przysiecki N, Seifert M, Keller MA, Oberhuber G, Sprung S, Ran Q, Koch R, Effenberger M, Tancevski I, Zoller H, Moschen AR, Weiss G, Becker C, Rosenstiel P, Kaser A, Tilg H, Adolph TE. Dietary lipids fuel GPX4-restricted enteritis resembling Crohn’s disease. Nature Communications. 2020;11(1):1775. doi: 10.1038/s41467-020-15646-6.

45. Aihaiti Y, Zheng H, Cai Y, Tuerhong X, Kaerman M, Wang F, Xu P. Exploration and validation of therapeutic molecules for rheumatoid arthritis based on ferroptosis-related genes. Life Sci. 2024;351:122780. Epub 20240610. doi: 10.1016/j.lfs.2024.122780. PubMed PMID: 38866217.

46. Tao K, Tian Y, Li S, Ni B, Song Z, Zhai Z. Ferroptosis in peripheral blood mononuclear cells of systemic lupus erythematosus. Clin Exp Rheumatol. 2024;42(3):651–7. Epub 20240111. doi: 10.55563/clinexprheumatol/kylvva. PubMed PMID: 38294021.

47. Wu K, Zhu E, Chen J, Kuang Q, Lin J, Zhao S, Xu X, Li S, Sui Y, Huang M, Zhang Y. Overexpression of GPX4 in diabetic rat kidney alleviates renal injury induced by ferroptosis. BioMetals. 2025;38(4):1281–97. doi: 10.1007/s10534-025-00706-5.

48. Klöditz K, Fadeel B. Three cell deaths and a funeral: macrophage clearance of cells undergoing distinct modes of cell death. Cell Death Discovery. 2019;5(1):65. doi: 10.1038/s41420-019-0146-x.

49. Doran AC, Yurdagul A, Jr., Tabas I. Efferocytosis in health and disease. Nat Rev Immunol. 2020;20(4):254–67. Epub 20191210. doi: 10.1038/s41577-019-0240-6. PubMed PMID: 31822793; PMCID: PMC7667664.

50. Zizzo G, Cohen PL. The PPAR-γ antagonist GW9662 elicits differentiation of M2c-like cells and upregulation of the MerTK/Gas6 axis: a key role for PPAR-γ in human macrophage polarization. Journal of Inflammation. 2015;12(1):36. doi: 10.1186/s12950-015-0081-4.

51. Michlewska S, Dransfield I, Megson IL, Rossi AG. Macrophage phagocytosis of apoptotic neutrophils is critically regulated by the opposing actions of pro-inflammatory and anti-inflammatory agents: key role for TNF-alpha. Faseb j. 2009;23(3):844–54. Epub 20081029. doi: 10.1096/fj.08-121228. PubMed PMID: 18971259.

52. McPhillips K, Janssen WJ, Ghosh M, Byrne A, Gardai S, Remigio L, Bratton DL, Kang JL, Henson P. TNF-alpha inhibits macrophage clearance of apoptotic cells via cytosolic phospholipase A2 and oxidant-dependent mechanisms. J Immunol. 2007;178(12):8117–26. doi: 10.4049/jimmunol.178.12.8117. PubMed PMID: 17548650.

53. Singh MV, Kotla S, Le N-T, Ae Ko K, Heo K-S, Wang Y, Fujii Y, Thi Vu H, McBeath E, Thomas TN, Jin Gi Y, Tao Y, Medina JL, Taunton J, Carson N, Dogra V, Doyley MM, Tyrell A, Lu W, Qiu X, Stirpe NE, Gates KJ, Hurley C, Fujiwara K, Maggirwar SB, Schifitto G, Abe J-i. Senescent Phenotype Induced by p90RSK-NRF2 Signaling Sensitizes Monocytes and Macrophages to Oxidative Stress in HIV-Positive Individuals. Circulation. 2019;139(9):1199–216. doi: doi:10.1161/CIRCULATIONAHA.118.036232.

54. Canton J, Khezri R, Glogauer M, Grinstein S. Contrasting phagosome pH regulation and maturation in human M1 and M2 macrophages. Mol Biol Cell. 2014;25(21):3330–41. Epub 20140827. doi: 10.1091/mbc.E14-05-0967. PubMed PMID: 25165138; PMCID: PMC4214780.

55. Bagaitkar J, Huang J, Zeng MY, Pech NK, Monlish DA, Perez-Zapata LJ, Miralda I, Schuettpelz LG, Dinauer MC. NADPH oxidase activation regulates apoptotic neutrophil clearance by murine macrophages. Blood. 2018;131(21):2367–78. doi: 10.1182/blood-2017-09-809004.

56. Viaud M, Ivanov S, Vujic N, Duta-Mare M, Aira LE, Barouillet T, Garcia E, Orange F, Dugail I, Hainault I, Stehlik C, Marchetti S, Boyer L, Guinamard R, Foufelle F, Bochem A, Hovingh KG, Thorp EB, Gautier EL, Kratky D, Dasilva-Jardine P, Yvan-Charvet L. Lysosomal Cholesterol Hydrolysis Couples Efferocytosis to Anti-Inflammatory Oxysterol Production. Circ Res. 2018;122(10):1369–84. Epub 20180309. doi: 10.1161/circresaha.117.312333. PubMed PMID: 29523554; PMCID: PMC6034181.

57. Cai S, Gao J, Weng X, Wang Z, Zheng D, Wang Q, Li Q, Han C, Li W, Chen J, Fu Y, Tan Y, Wei B, Pang Z, Huang Z, Song Y, Ge J. Synergistic enhancement of efferocytosis and cholesterol efflux via macrophage biomimetic nanoparticle to attenuate atherosclerosis progression. Bioact Mater. 2026;55:131–43. Epub 20250919. doi: 10.1016/j.bioactmat.2025.09.022. PubMed PMID: 41035425; PMCID: PMC12481714.

58. Wu Y, Zhou H, Liu H, Hu J, Sun Y, Yan W, Tong C, Kong Y, Liu B. Pitavastatin-loaded procyanidins self-assembled nanoparticles alleviate advanced atherosclerosis via modulating macrophage efferocytosis and cholesterol efflux. Acta Pharm Sin B. 2025;15(6):3305–20. Epub 20250506. doi: 10.1016/j.apsb.2024.08.006. PubMed PMID: 40654339; PMCID: PMC12254700.

59. Boucher DM, Robichaud S, Lorant V, Leon JS, Suliman I, Rasheed A, Susser LI, Emerton C, Geoffrion M, De Jong E, Bowdish DME, Aikawa M, Aikawa E, Singh SA, Rayner KJ, Ouimet M. Age-Related Impairments in Immune Cell Efferocytosis and Autophagy Hinder Atherosclerosis Regression. Arterioscler Thromb Vasc Biol. 2025;45(4):481–95. Epub 20250213. doi: 10.1161/atvbaha.124.321662. PubMed PMID: 39945065; PMCID: PMC11936474.

60. Yildirim Z, Baboo S, Hamid SM, Dogan AE, Tufanli O, Robichaud S, Emerton C, Diedrich JK, Vatandaslar H, Nikolos F, Gu Y, Iwawaki T, Tarling E, Ouimet M, Nelson DL, Yates JR, 3rd, Walter P, Erbay E. Intercepting IRE1 kinase-FMRP signaling prevents atherosclerosis progression. EMBO Mol Med. 2022;14(4):e15344. Epub 20220222. doi: 10.15252/emmm.202115344. PubMed PMID: 35191199; PMCID: PMC8988208.

61. Chen W, Li L, Wang J, Zhang R, Zhang T, Wu Y, Wang S, Xing D. The ABCA1-efferocytosis axis: A new strategy to protect against atherosclerosis. Clin Chim Acta. 2021;518:1–8. Epub 20210316. doi: 10.1016/j.cca.2021.02.025. PubMed PMID: 33741356.

